# Multi-scale Imaging Reveals Aberrant Connectome Organization and Elevated Dorsal Striatal *Arc* Expression in Advanced Age

**DOI:** 10.1101/434191

**Authors:** Luis M. Colon-Perez, Sean M. Turner, Katelyn N. Lubke, Marcelo Febo, Sara N. Burke

## Abstract

The functional connectome reflects a network architecture enabling adaptive behavior that becomes vulnerable in advanced age. The cellular mechanisms that contribute to altered functional connectivity in old age, however, are not known. Here we used a multi-scale imaging approach to link age-related changes in the functional connectome to altered expression of the activity-dependent immediate-early gene *Arc* as a function of training to multi-task. Aged behaviorally-impaired, but not young, rats had a subnetwork of increased connectivity between the anterior cingulate cortex and dorsal striatum. Moreover, the old rats had less stable rich club participation that increased with cognitive training. The altered functional connectome of aged rats was associated with a greater engagement of neurons in the dorsal striatum during cognitive multi-tasking. These findings point to aberrant large-scale functional connectivity in aged animals that is associated with altered cellular activity patterns within individual brain regions.

---

Advancing age is associated with cognitive impairments that can erode one’s quality of life, even in the absence of pathology (Burke and Barnes, 2006; Samson and Barnes, 2013; Lockhart and DeCarli, 2014). Behaviors that rely on largescale interactions across brain networks, such as episodic memory or cognitive multitasking, appear to be particularly vulnerable to decline in older humans (Dennis et al., 2008; Chadick et al., 2014; Fandakova et al., 2014; Salami et al., 2014) and animal models of cognitive aging (Hernandez et al., 2015). A possible reason for these cognitive impairments in older adults could be aberrant connectome connectivity and organization with advancing age (Ash and Rapp, 2014; Ash et al., 2016). A powerful tool for evaluating the integrity of network organization is to quantify resting state functional connectivity and connectomics (Sala-Llonch et al., 2015a; Nyberg, 2017). In fact, several studies have reported that older adults have decreased functional connectivity within the default mode network (Sala-Llonch et al., 2015a; Grady et al., 2016), as well as increased functional connectivity within the hippocampal network (Salami et al., 2014), the frontoparietal control and the dorsal attentional networks (Grady et al., 2016), as well as between the anterior cingulate cortex and other cortical structures (Cao et al., 2014b) that relate to cognitive performance. Importantly, altered network functional connectivity has also been reported for old rats (Ash et al., 2016), indicating that there is a cross-species consensus regarding the vulnerability of these networks to advancing age.

While altered network connectivity in older adults is thought to reflect neural inefficiency or a dedifferentiation process that is associated with cognitive decline (Salami et al., 2014; Sala-Llonch et al., 2015b; Grady et al., 2016; Nyberg, 2017), it remains unclear how network parameters used to quantify the large-scale functional connectome organization relate to age-associated neurobiological alterations at the cellular level. Recent behavioral models for probing the integrity of inter-regional communication (Hernandez et al., 2015; Hernandez et al., 2017), along with advances in small animal functional MRI (Ash et al., 2016; Colon-Perez et al., 2016) offer a unique opportunity to study the brain’s connectome organization using functional connectivity in conjunction with quantification of the neurobiological variables enabling scales from large networks to individual neurons to be bridged within the same animals. Critically, altered functional connectivity in rat models is also predictive of cognitive decline in advanced age (Ash et al., 2016), indicating that this experimental system can be used to link large-scale network decline to specific neurobiological alterations.

An additional advantage to working with animal models is the ability to longitudinally measure resting state metrics of network architecture as a function of cognitive training in populations with highly controlled dietary and behavioral experiences across age groups.

Previous longitudinal studies have shown that resting state brain networks are stable over time (Iordan et al., 2017). Importantly, however, working memory training in young adults can elicit plasticity in network architecture (Takeuchi et al., 2017). Little is known, however, regarding the ability of cognitive training to alter functional network architecture in aged populations, and if this is comparable to what is observed in younger study participants.

The current study aimed to examine how cognitive training on a cognitive dual task, which required rats to perform a spatial working memory and a biconditional association task (WM/BAT) simultaneously, altered resting-state functional connectivity in young and aged rats. This behavioral paradigm is known to require interactions between prefrontal, medial temporal and subcortical structures (Jo and Lee, 2010; Hernandez et al., 2017), and is vulnerable to decline in old age prior to the emergence of deficits on the hippocampus-dependent Morris water maze (Hernandez et al., 2015). Rats were scanned at three distinct time points enabling the determination of longitudinal changes in brain connectivity prior to and during learning. Following the last resting state scan, rats were retrained on a novel WM/BAT problem set to allow for the assessment of expression of the activity-dependent immediate-early gene *Arc* (Cole et al., 1989; Guzowski et al., 1999) to relate direct neuronal activity during the task to network connectivity patterns obtained from fMRI resting state data. Thus, the current experiments used a multi-scale imaging approach that spanned cells to global networks, integrating connectomics, *Arc* expression patterns, and behavior to explore the changes in brain connectivity in young and old rats during cognitive training.

## Results

### Working memory/biconditional association task (WM/BAT) performance

Rats were placed on food restriction, and then initially shaped to traverse a digital-8-shaped maze for ‘Froot Loop’ rewards (The Kellogg Company, Battle Creek, MI, USA). The working memory/biconditional association task (WM/BAT; Figure 1a) required animals to alternate between making left and right turns on a digital-8-shaped maze. A choice platform with 2 food wells was located on each side of the maze. When an animal reached the choice platform, after correctly alternating, they were presented with two objects (e.g., a dog and an owl figurine) placed over each food well. On the left choice platform, the dog was the rewarded object and animals received a foot loop piece for displacing it. On the right choice platform, the owl was the correct object selection. Thus, animals had to multi-task by acquiring an object-in-place rule while simultaneously performing continuous spatial alternations. Resting state functional MRI scans were obtained following food restriction and 1 week of pretraining to run on the maze. This baseline scan was acquired within 2 days of the last pre-training session and prior to the initiation of WM/BAT training. A second and third resting scan was obtained after 11 (scan 2) and 27 days (scan 3) of training, respectively. Starting the pretraining maze exposure and food restriction prior to imaging ensured that changes in functional connectivity were not induced by introducing these behavioral conditioning procedures between imaging sessions. Following the third scan, rats were trained on a new WM/BAT problem set with a unique pair of objects for the *Arc* experiment.

**Figure 1:**
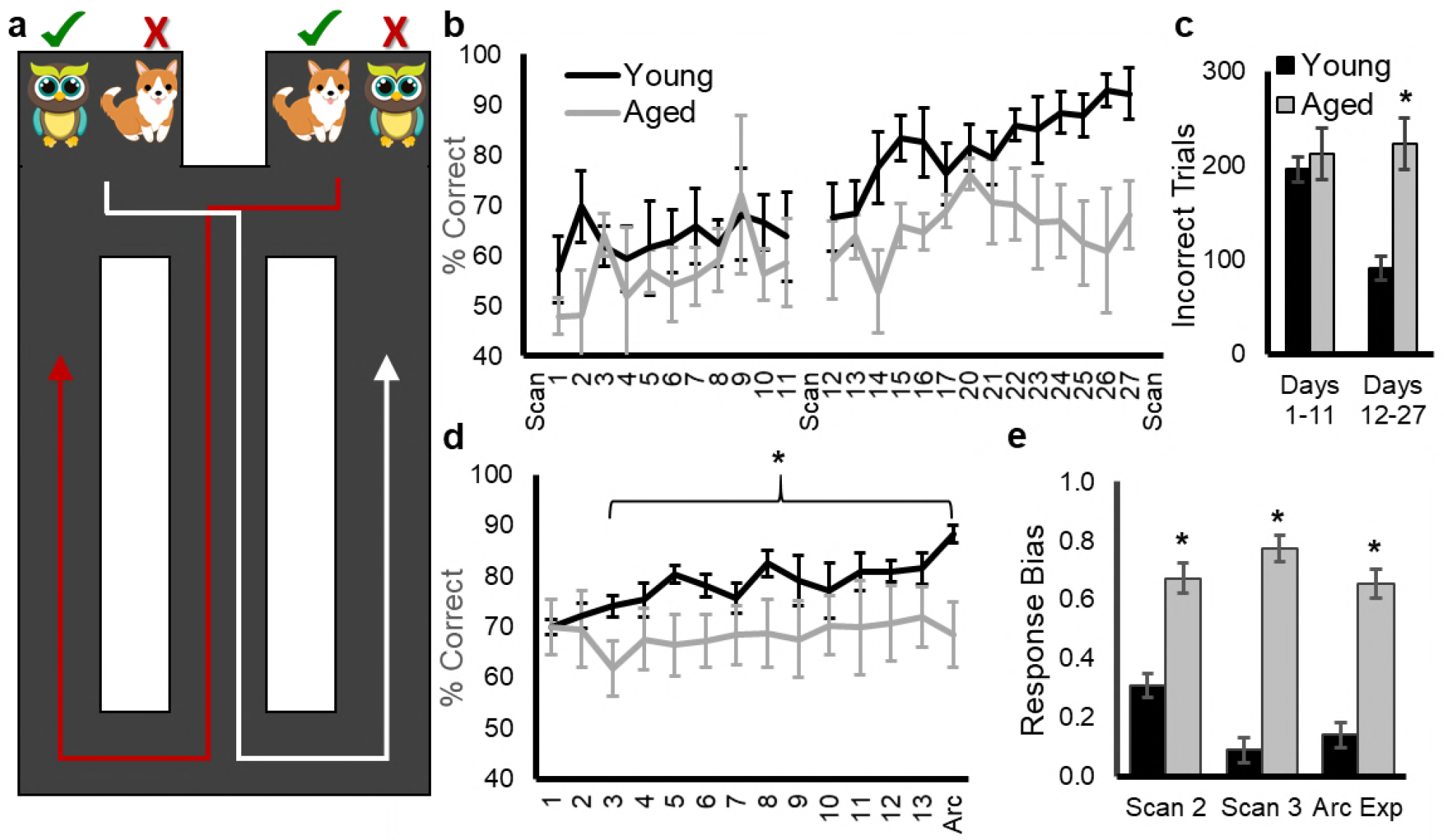
Working memory/biconditional association task (WM/BAT) performance. **(a)** Schematic of the WM/BAT. Rats traversed a figure-8-shaped maze alternating between left and right turns. After turning, before returning to the central stem, rats solved an object discrimination problem in which the correct choice (green check) was contingent on side of the maze. On the left, the owl is rewarded and on the right the dog is rewarded. **(b)** Performance on WM/BAT over days of testing task in young (black) and aged (grey) rats between resting state scans. **(c)** The total number of incorrect trials made between scans 1 and 2 (Days 1-11) and scans 2 and 3 (Days 12-27). Aged rats made significantly more errors across testing days 12-27. **(d)** After the third scan, rats were re-trained on WM/BAT with different objects for 13 days before performing the task a final time, followed by immediate sacrifice to label the mRNA products of the activity-dependent immediate-early gene Arc. **(e)** The response bias of young and aged rats on the days prior to scans 2 and 3 and on the day of the Arc experiment. Aged rats had a significantly greater response bias compared to young rats across all time points. Error bars are ± 1 SEM, *p < 0.05.

Figure 1b-d summarizes the mean performances of young (black) and aged (grey) rats across testing days for the first (Figure 1b/c) and the second (Figure 1d) WM/BAT problem sets. Between the first and second scan, there was not a significant main effect of testing day (F_[10,80]_ = 0.74, p = 0.68), age (F_[1,8]_ = 1.71, p = 0.27), or an age by test day interaction (F_[10,80]_ = 0.45, p = 0.92). In contrast, between the second and third scan, there was a significant main effect of testing day (F_[13,91]_ = 3.84, p = 0.019). Orthogonal contrasts comparing each day of testing to performance on the day following the second scan (Day 12) indicated that the percentages of correct responses were significantly greater by day 17 compared to day 12 (p = 0.038). There was also a trend for an age effect (F_[1,8]_ = 4.30, p = 0.07), but no significant interaction between age and test day (F_[13,91]_ = 1.09, p = 0.38). Another way to evaluate the performances of young and aged rats is to compare the total number of incorrect trials during training (Figure 1c). Across all days of testing, the aged rats made significantly more errors than the young rats (F_[1,8]_ = 10.02, p = 0.013). Importantly, there was a significant interaction effect between age and phase of testing (Days 1-11 versus Days 12-27; F_[1,8]_ = 5.54, p = 0.046). Specifically, post hoc analysis indicated that young and aged rats made a similar number of errors prior to the second scan (Days 1-11; T_[8]_ = 0.55, p = 0.60), but aged rats made significantly more errors prior to the third scan (Days 12-27; T_[8]_ = 3.50, p = 0.008, corrected α = 0.025).

After the third scan, rats were retrained on WM/BAT with different objects for 13 days before performing the task a final time, followed by immediate sacrifice to label the mRNA products of the activity-dependent immediate-early gene *Arc (Guzowski et al., 1999)*. During re-training, there was a significant main effect of testing day (F_[13,91]_ = 2.91, p < 0.01), with rats showing improvements across days. There was also a significant interaction effect between age and test day (F_[13,91]_ = 2.14, p < 0.02), with aged rats performing similar to young on the first two test days, but significantly worse on the final day (F_[1,7]_ = 5.29, p < 0.05).

Previous studies have reported that before animals learn an object discrimination problem, they show a significant response bias by selecting an object over a food well on a particular side (left versus right) regardless of object identity (Lee and Byeon, 2014; Hernandez et al., 2015; Johnson et al., 2017a). This innate response bias must be overcome before animals will learn the biconditional rule (Lee and Byeon, 2014). The response bias was calculated for young and aged rats on the days before scans 2 and 3 and on the day of the *Arc* experiment (Figure 1e). There was not a significant effect of day on the response bias (F_[2,14]_ = 1.42, p = 0.27), but aged rats had a significantly larger response bias relative to the young animals (F_[1,7]_ = 124.19, p = 0.0001), consistent with previous reports (Hernandez et al., 2015; Johnson et al., 2017a). The interaction effect between age and day was not significant (F_[2,14]_ = 1.32, p = 0.30), indicating that the elevated response bias of the aged rats persisted throughout testing.

### Resting state connectivity

Over the past decade, graph theoretical approaches have been widely used to quantify functional brain networks (Bullmore and Sporns, 2009, 2012; Ash and Rapp, 2014). This analytical approach models the brain as a complex network composed of nodes (i.e., brain regions) and edges (i.e.,functional correlations) connecting the nodes (Bullmore and Sporns, 2009). Figure 2a/b shows the 3D brain networks with functional edges with z scores larger than 0.3 in young and aged rats across scanning sessions. Metrics for global network connectivity (e.g., node strength, degree, weighted and binary path lengths and clustering coefficients) were not significantly affected by scanning session, age, nor did the scanning session by age interaction reach significance (see Table 1 for statistical summary). These findings are consistent with previous data that resting state networks are consistent over time (Iordan et al., 2017).

**Table 1:** Quantification of global inter-node connectivity patterns.

| Variable | Scanning Session | Age | Scanning Session x Age Interaction |
| --- | --- | --- | --- |
| Node degree | $F_{[2,29]} = 0.50$, $p = 0.61$ | $F_{[1,29]} = 2.00$, $p = 0.17$ | $F_{[2,29]} = 0.50$, $p = 0.61$ |
| Node strength | $F_{[2,29]} = 1.06$, $p = 0.36$ | $F_{[1,29]} = 0.52$, $p = 0.47$ | $F_{[2,29]} = 0.21$, $p = 0.81$ |
| Path length | $F_{[2,29]} = 0.86$, $p = 0.44$ | $F_{[1,29]} = 0.59$, $p = 0.45$ | $F_{[2,29]} = 0.10$, $p = 0.90$ |
| Clustering coefficient | $F_{[2,29]} = 0.54$, $p = 0.59$ | $F_{[1,29]} = 0.11$, $p = 0.73$ | $F_{[2,29]} = 0.41$, $p = 0.67$ |

**Figure 2:**
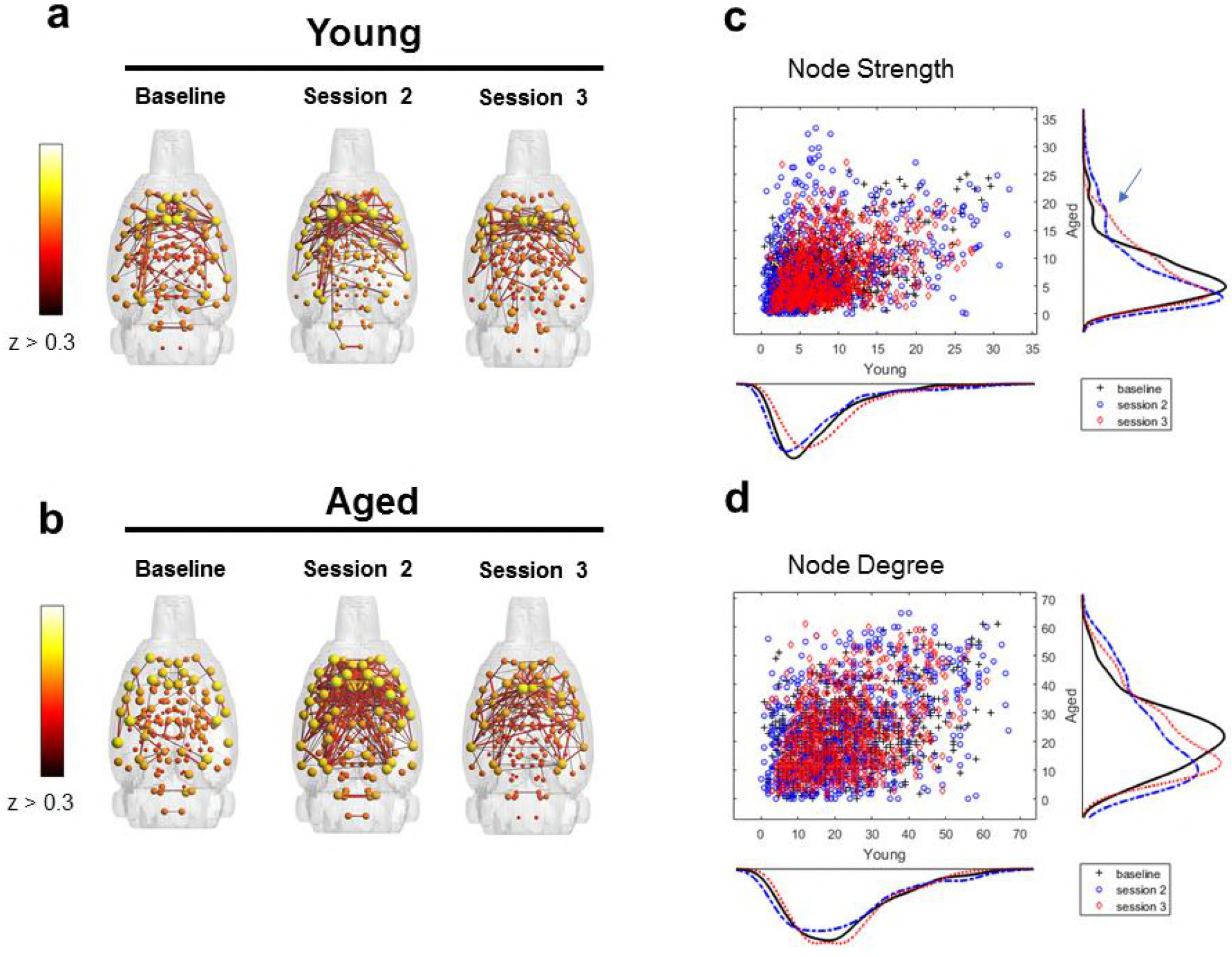
Connectivity patterns by scanning session and age group. Connected modules with edges z > 0.3 for young **(a)** and aged **(b)** rats across scan sessions. Connectivity indices indicate larger network engagement between baseline and the second and third scanning sessions in aged (blue arrow), but not young rats. This is indicated by more nodes with node strength > 15 **(c)** and degree > 40 **(d)**.

The node strength and node degree distributions, however, displayed a local increase of nodes with high strengths (s > 15; Figure 2c) and degree (k > 40; Figure 2d) in the aged cohort between the baseline and subsequent scan sessions (blue arrows; Figure 2c/d). From this distribution, we identified the nodes with strength values larger than 15 during the second scan session in the aged rats, 16 nodes in total. Figure 3 shows the connectivity between these nodes in young (Figure 3a) and aged (Figure 3b) rats across scanning session, as well as the associated brain regions (Figure 3c). Network parameters for this subnetwork showing increased connectivity following cognitive training was compared between young and aged rats with a factorial ANOVA with scan session used as a within-subject factor. There was a significant main effect of scan session on the q parameter of modularity (F_[2,29]_ = 4.42, p = 0.02), indicating increased modularity after cognitive training. Modularity did not significantly change as a function of age (F_[1,29]_ = 0.01, p = 0.94), nor was the interaction between age and scan session significant (F_[2,29]_ = 0.17, p = 0.20). Similar to modularity, there was a main effect of scan session on the clustering coefficient (F_[2,29]_ = 3.39, p = 0.05), but no effect of age (F_[1,29]_ = 0.05, p = 0.82) or a scan session x age interaction (F_[2,29]_ = 0.39, p = 0.68). Finally, the node strength showed a significant main effect of scanning session (F_[2,29]_ = 3.85, p = 0.04), but not of age (F_[1,29]_ = 0.02, p = 0.90) or a scan session × age interaction (F_[2,29]_ = 0.30, p = 0.74). These data indicate that while the global network parameters did not change with scan session, the high degree nodes become more modular and nodal interactions are stronger in both the aged and young rats.

**Figure 3:**
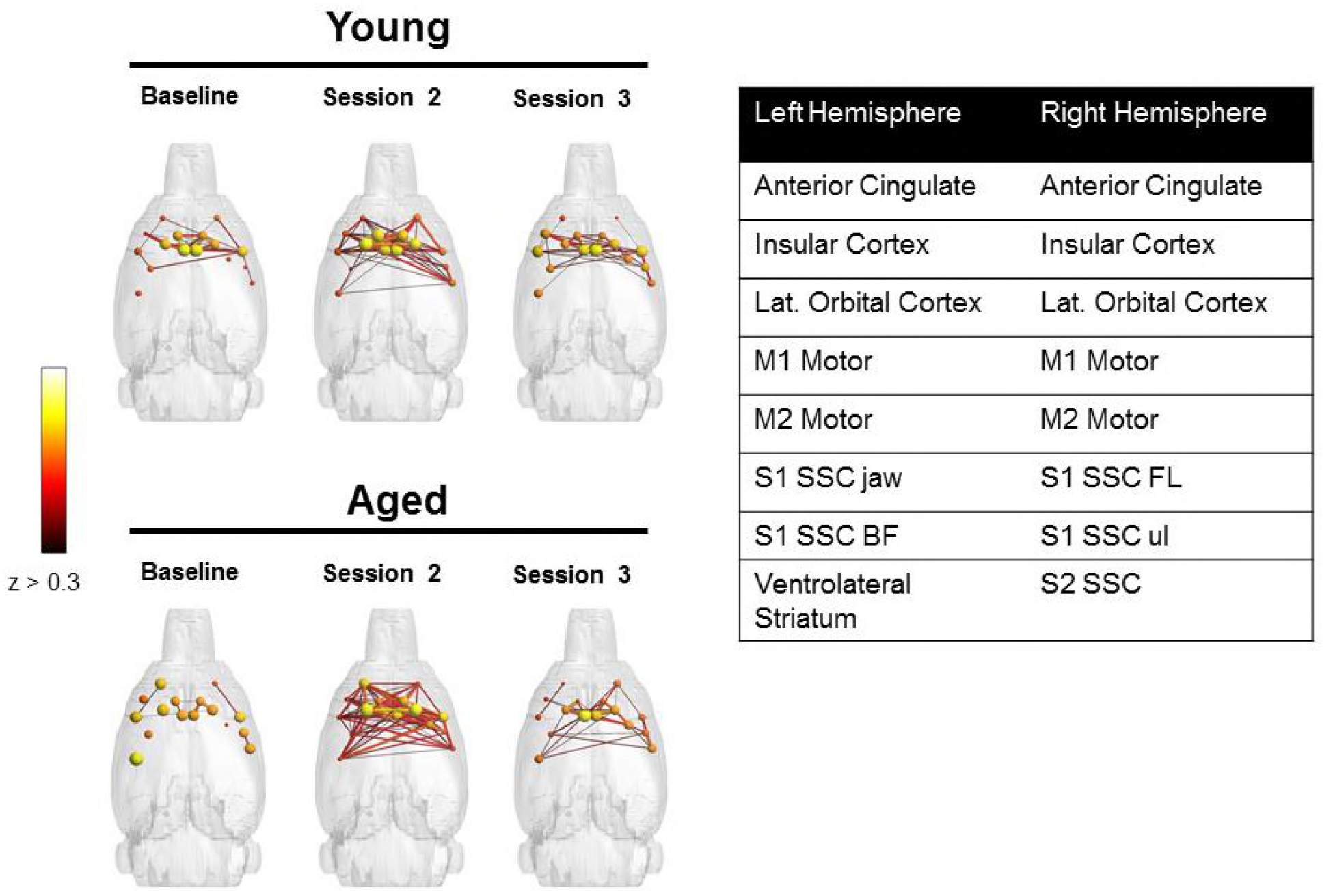
Nodes with increased strength. 16 nodes with strength > 15 during scan 2 in aged rats were identified from the distribution shown in Figure 1. Connectivity patterns of these nodes for the young **(a)** and aged **(b)** groups. The table in **(c)** lists the regions by hemisphere that correspond to these nodes.

Because the anterior cingulate cortex (ACC) was identified as a node with higherstrength values in both hemispheres, and there are known age-related physiological changes within this region (Insel et al., 2012; Insel and Barnes, 2015), and morphological differences in this structure are implicated in successful aging (Rogalski et al., 2012), we used the ACC as a seed to quantify functional connectivity between this region and other nodes as a function of scan session (Figure 4a). Using this approach, we observed that functional connectivity between ACC and dorsal striatum (DS) changed as a function of scan session and age (Figure 4b). Although the main effect of age (F_[1,8]_ = 0.72; p = 0.42) and scan session (F_[2,16]_ = 0.56; p = 0.58) did not reach statistical significance, the interaction between age and scan session was significant (F_[2,16]_ = 6.75; p = 0.008). Post hoc analysis indicated that there were no significant age differences between ACC-DS connectivity during the baseline scan (95% confidence interval: −0.25 to 0.67, p = 0.57), and scan 2 (95% confidence interval: −0.49 to 0.43, p = 0.99), but the aged rats had significantly higher connectivity relative to young during the third scan (95% confidence interval: −0.99 to −0.07, p = 0.02). These data are interesting in the context of the behavioral results showing that aged rats have a significantly larger response bias across training relative to the young (Figure 1e). It is well established that the DS is involved in response-based learning strategies (Packard and McGaugh, 1992; Gold, 2004), and aged rats may default to more response-based strategies as spatial learning becomes impaired (Tomas Pereira et al., 2015). Thus, the increased ACC-DS connectivity observed here may be a network signature of the enhanced response bias seen at the behavioral level.

**Figure 4:**
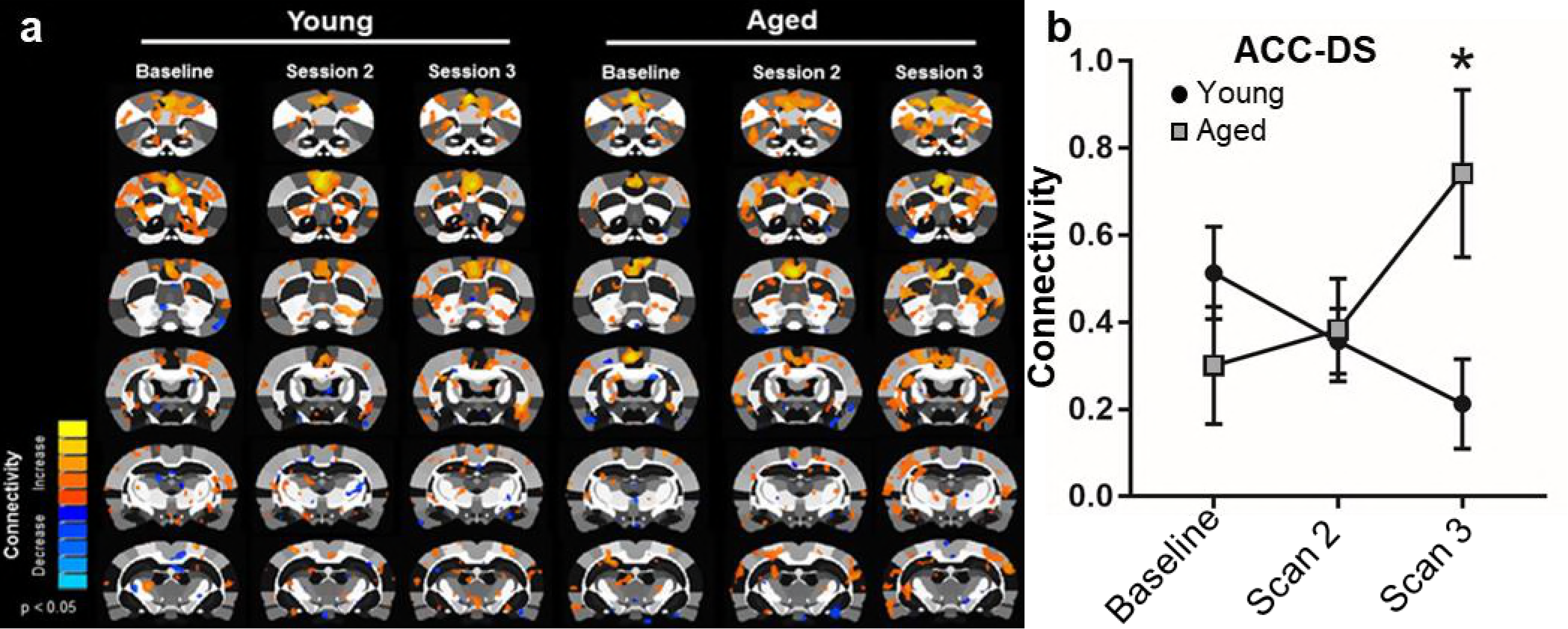
Seed analysis of ACC connectivity. **(a)** Connectivity with ACC in young (left) and aged (right) rats as a function of scan session. **(b)** Connectivity between the ACC and DS as a function of scan session in young (black) and aged (grey) rats. The main effect of age (F_[1,8]_ = 0.72; p = 0.42) and scan session (F_[2,16]_ = 0.56; p = 0.58) did not reach statistical significance. The interaction effect between age and scan session was significant, however (F_[2,16]_ = 6.75; p = 0.008). Post hoc analysis indicated that there were no significant age differences between ACC-DS connectivity during the baseline scan (p = 0.57), and scan 2 (p = 0.99), but the aged rats had significantly higher connectivity relative to young during the third scan (p = 0.02).

Because of the potential presence of a subnetwork of ‘hub’ nodes driving the highly interconnected and modular network patterns in older rats, we next assessed the “rich club” index. The rich club refers to a set of densely and highly inter-connected nodes known as hub regions in the brain. Rich-club organization is an expensive network structure (i.e., extensive connectivity and metabolic cost) that allows complex network dynamics to increase brain function efficiency (Kaiser and Hilgetag, 2006; van den Heuvel et al., 2012). The structural rich club (brain regions connected by white matter tracts) is described as a connectivity backbone allowing an efficient information transfer between distant brain regions (van den Heuvel et al., 2012). In the context of functional networks (brain regions connected by correlations derived from fMRI), the rich club describes an increase in participation and activation of certain active nodes into members of a functional rich club of otherwise structurally weakly connected nodes (Liang et al., 2018). Thus, rich club indices were calculated for young and aged rats as a function of scanning session. Figure 5 shows the rich club organization in young and aged rats at baseline (Figure 5a) and after cognitive training as a function of scan session (Figures 5 b/c). Similar to previous reports from human study participants (Cao et al., 2014a), the functional rich club architecture at the baseline scan was significantly reduced in aged rats relative to young for rich club indices (k-level) > 27 (p < 0.05). The young rats had stable functional rich club architecture across all scans (Figure 5b), which has previously been reported (Liang et al., 2018). Interestingly, in the aged rats there was a significant increase in rich club participation after the baseline scan (Figure 5c). Starting after the first training period (11 days), rich club for indices larger than 20 displayed a significant main effect of scanning session (F_[2,29]_ > 3.4, p < 0.05), but not of age (F_[1,29]_ < 0.55, p > 0.5). The interaction term of the rich club displayed an interaction term for large k levels (k > 28) (F_[2,29]_ > 3.80, p < 0.05). As evident in Figures 5b/c, the significant interaction between scan session and rich club participation was due to the aged rats having greater rich club indices after cognitive training, while the young rats did not show a change. These changes were specific to rich club participation since no global changes in topological indices of small-worldness, and modularity indices were observed (see supplemental data).

**Figure 5.**
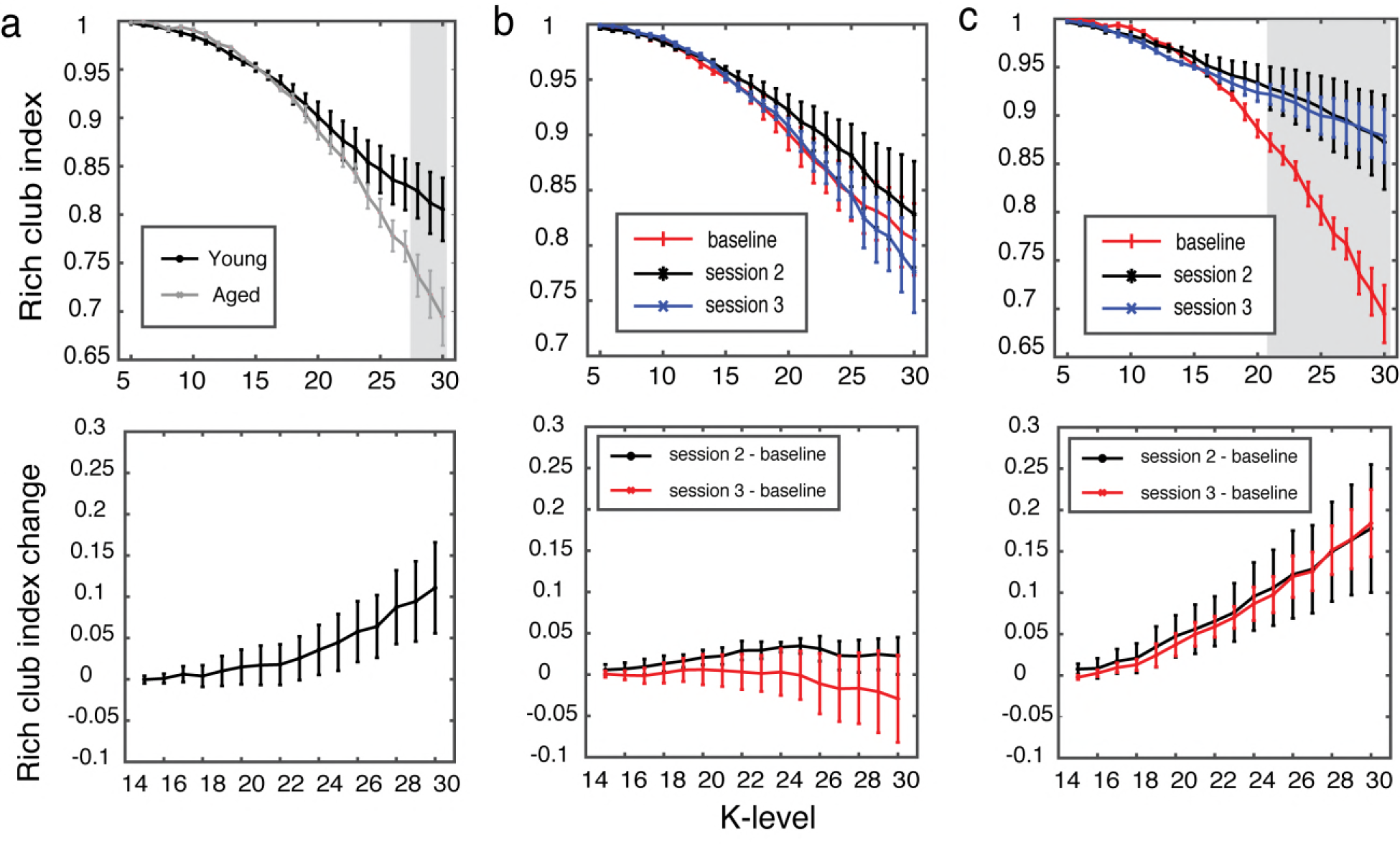
Rich club organization increases with cognitive training in aged rats. **(a)** Top figure, rich club participation index between young and aged rats, at the baseline scan (shaded area K > 27 and p < 0.05), bottom figure shows the overall difference increase for between aged and young rats. **(b)** Top figure, young rats rich club participation index, and bottom is the difference between scanning sessions. No changes in any of the sessions or any K-level was observed. **(c)** Top figure, aged rats rich club participation in aged group increase following cognitive training for large K-levels (shaded area K > 21 and p < 0.05), bottom figure shows the overall difference increase for both sessions.

### Behaviorally-induced expression of the immediate-early gene Arc

The resting state data identified age-related differences in global connectivity patterns as a function of cognitive training. These data, however, cannot provide information regarding cellular activity patterns during behavior. Thus, to examine potential age-related differences in neuron activity during WM/BAT performance, we trained young and aged rats on a new problem set for 13 days. On the 14^th^ day of testing, rats performed either the WM/BAT or a control spatial alternation task in which a food reward was randomly placed in a food well on the choice platform. The reward was not covered and rats did not have to perform an object discrimination in this control task. After this first epoch of behavior, rats were placed in their home cages for a 20-min rest. Following the rest, rats performed a second epoch of behavior for 5 min. All rats performed one epoch of WM/BAT and one epoch of spatial alternation in counterbalanced order. The performances of young and aged rats on WM/BAT is shown in Figures 1d/e. No rats made errors on the control alternation task.

Immediately, after the second epoch, rats were heavily sedated in an anesthesia chamber with concentrated isoflurane and decapitated. Brains were rapidly extracted, and tissue was processed to label the mRNA products of the immediate-early gene *Arc* for cellular compartment analysis of temporal activity with fluorescence *in situ* hybridization (catFISH). The subcellular localization of *Arc* mRNA can be used to determine which neuronal ensembles across the brain were active during 2 distinct episodes of behavior. *Arc* is first transcribed within the nucleus of neurons 1-2 minutes after cell firing. Importantly, *Arc* mRNA translocate to the cytoplasm approximately 15-20 minutes after cell firing, which allows for cellular activity during 2 epochs of behavior, separated by a 20-min rest to be represented within a single neural population (Guzowski et al., 1999).

Due to the elevated response bias of aged rats (Figure 1e), and the observation that the old animals showed an increase in ACC-DS connectivity as a function of cognitive training, we focused our *Arc* catFISH analysis on the ACC and DS. On the day of the catFISH experiment, repeated-measures ANOVA with the within subject factor of task and the between subjects factor of age group did not show a significant difference in the number of WM/BAT and spatial alternation trials completed (F_[1,7]_ = 2.31, p = 0.11). Moreover, the main effect of age did not reach statistical significance (F_[1,7]_ = 3.31, p = 0.11), nor was the interaction between age and task significant (F_[1,7]_ = 0.61, p = 0.46). Thus, any differences in neuron activation could not be explained by animals performing a different number of trials.

Figure 6a shows the region of ACC that was imaged and representative examples of *Arc* labeling in a young and an aged rat. The percentage of ACC neurons activation during the WM/BAT and spatial alternation task are shown for young and aged rats in Figure 6b. Repeated-measures ANOVA with the within subject factor of task and the between subjects factors of age group, hemisphere and cortical layer did not detect a significant difference in the proportion of cells activated during WM/BAT versus spatial alternation (F_[1,28]_ = 1.53, p = 0.23). Additionally, none of the other between subjects factors reached statistical significance (F_[2,28]_ < 3.01, p > 0.09, for all comparisons), nor were any of the interaction terms significant (F_[1,28]_ < 2.99, p > 0.1, for all comparisons).

**Figure 6:**
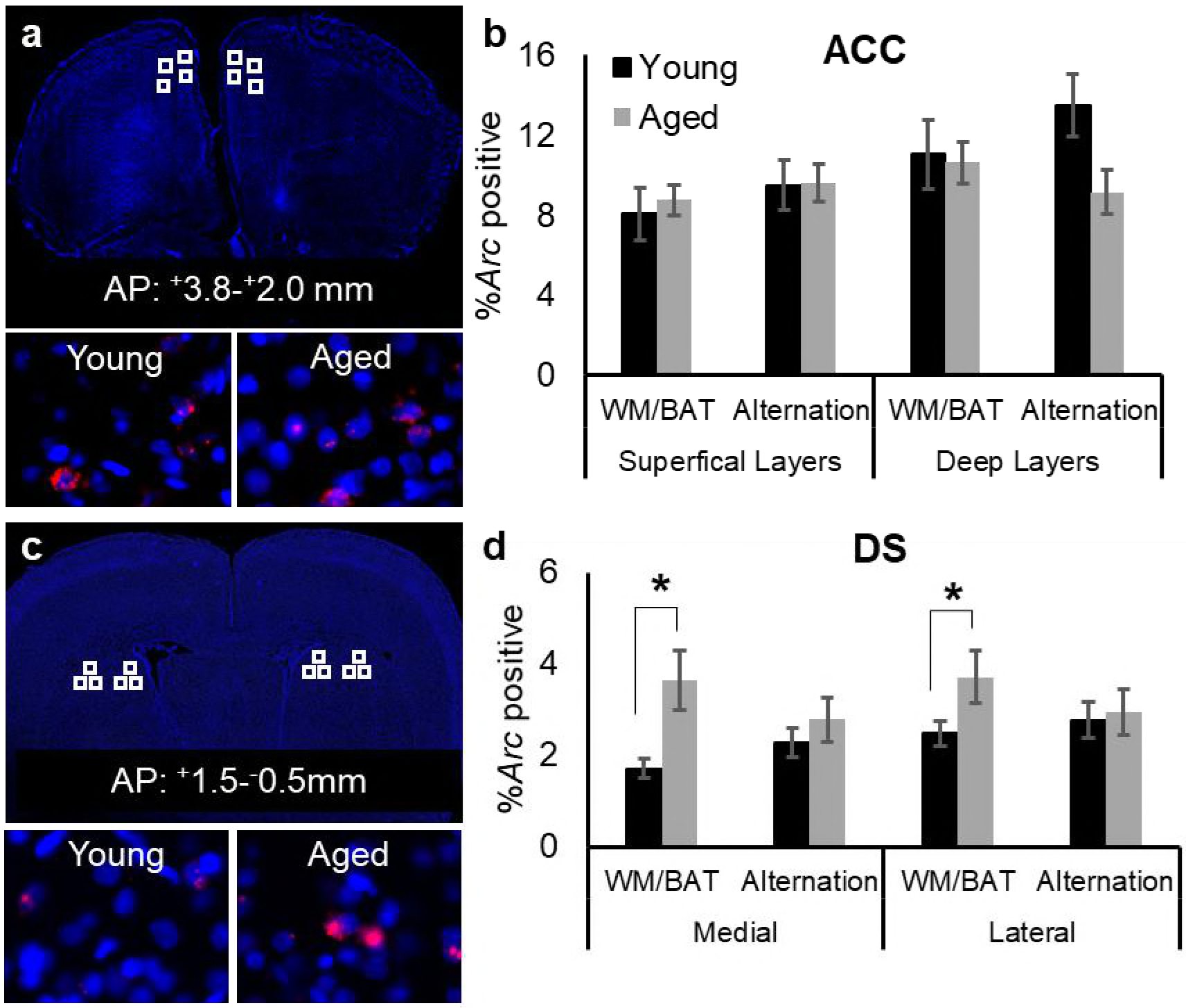
Neuronal activation during WM/BAT and spatial alternation in young and aged rats. **(a)** A DAPI stained section showing regions that were sampled (white squares) within the anterior cingulate cortex (ACC). Bottom panels show representative *Arc* labeling in a young (left) and an aged **B.** Representative images labeled for *Arc* mRNA (red) and DAPI (cell bodies; blue) from a young (top) and an aged (bottom) rat. **C.** Percentage of *Arc* positive cells corresponding to transcription during the bi-conditional association task, or the alternation task in young (black) and ages (grey) rats. There was not a significant main effect of task (F_[1,28]_ = 1.53, p = 0.22), or age (F_[1,28]_ = 0.68, p = 0.42) on the percentage of cells positive for *Arc*. Moreover, none of the interaction effects reached statistical significance (p > 0.1 for all comparisons).

Figure 6c shows the areas of the DS that images were collected from, as well as representative examples of *Arc* labeling in young and aged rats. Samples were taken from both the medial and lateral DS. Figure 5d shows the percentages of neurons that were positive for *Arc* during the different tasks in young and aged rats. Repeated-measures ANOVA with the within subject factor of task and the between subjects factors of age group, hemisphere and subregion (medial versus lateral DS) indicated that there was not a significant main effect of task on the percentage of cells positive for *Arc* (F_[1,28]_ = 1.24, p = 0.28). The aged rats, however, had significantly more cells that were positive for *Arc* compared to the young animals (F_[1,28]_ = 5.58, p < 0.03). This age difference was observed in both the medial and lateral DS, as indicated by lack of a main effect of subregion (F_[1,28]_ = 0.85, p = 0.37). The interaction effect between age and task was also significant (F_[1,28]_ = 13.64, p < 0.001), such that aged rats had more cells than young rats that transcribed *Arc* during WM/BAT (p < 0.001), but this same difference was not observed during the alternation task (p = 0.43). These data indicate that the enhanced DS activation in aged rats was specific to the behavioral in task in which the old animals demonstrated a deleterious response bias associated with worse performance. No other interaction effects reached statistical significance (F_[1,28]_ < 1.81, p > 0.18, for all comparisons).

To examine the extent that the active neuronal ensemble changed between the different tasks, a similarity score was calculated. As population overlap can be affected by differences in overall activity levels, similarity scores can be calculated to control for differences in activity between regions (Vazdarjanova and Guzowski, 2004). The similarity scores were compared between the ACC and DS (within subjects factor of region), and the between subjects variables of age and hemisphere. ‘Since layer, subregion and hemisphere did not have a significant effect activation within the ACC and DS, these factors were left out of the similarity score analysis. Figure 7 shows the average similarity scores for young and aged rats. The main effect of region was not significant (F_[1,34]_ = 0.01, p = 0.92), indicating that the ACC and DS updated activity patterns similarly in response to performing a different task within the same environment. Interestingly, there was a significant main effect of age on similarity score (F_[1,34]_ = 6.70, p < 0.02), but the interaction effect between region and age did not reach statistical significance (F_[1,34]_ = 0.11, p = 0.74). Together these data indicate that the aged rats had less population overlap across tasks compared to the young rats in both the ACC and DS. The reduced overlap in aged rats would reflect the difference between using a response-based strategy during WM/BAT that was not utilized during the spatial alternation task.

**Figure 7:**
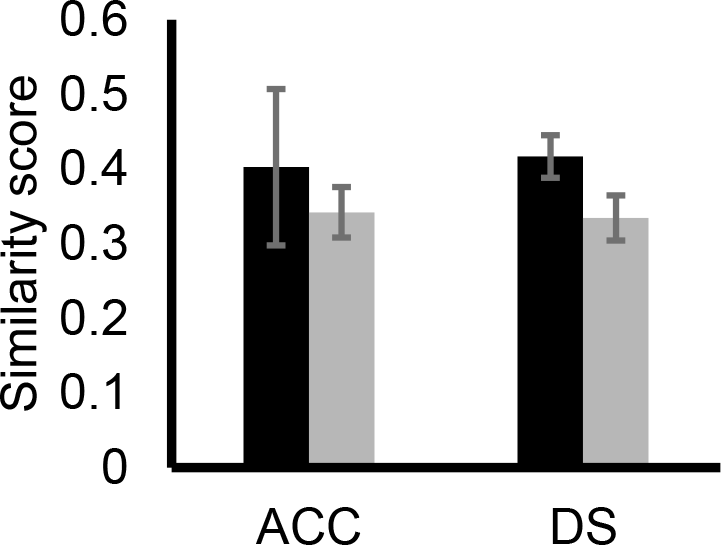
Population overlap in ACC and DS. The average similarity score for young (black) and aged (grey) rats in the ACC and the DS. The main effect of region was not significant (F_[1,34]_ = 0.01, p = 0.92). There was a significant main effect of age on similarity score (F_[1,34]_ = 6.70, p < 0.02), but the interaction effect between region and age did not reach statistical significance (F_[1,34]_ = 0.11, p = 0.74).

## Discussion

The current study used a multi-scale imaging approach, integrating resting state fMRI data with single-cell imaging of neuron activity, to determine the global network changes and cellular activity patterns in young and aged rats in relation to cognitive training. Critically, the resting state data were acquired longitudinally as a function of training on a working memory/bi-conditional association task (WM/BAT). Similar to a previous report, the aged rats were impaired on the WM/BAT relative to the young animals (Hernandez et al., 2015). Additionally, the aged rats showed an aberrant response bias during testing such that they often selected an object on one side of the choice platform, regardless of their location on the maze (left versus right platform), or the object identity (Figure 1e). This response bias in aged rats has been reported in other behavioral experiments (Hernandez et al., 2015; Johnson et al., 2017b). Importantly, the ability to inhibit this response-driven strategy is dependent on the medial prefrontal cortex and is associated with performance improvements (Lee and Byeon, 2014).

Several novel findings were found in this study. First, aged rats displayed an increase in brain connectivity among high degree nodes between the first baseline scan session and the subsequent sessions after training. This increase in connectivity among the high degree nodes, occurred independent of any significant increases in cognitive performance (Figures 2 and 3). The increase in connectivity was evident in the anterior cingulate cortex (ACC) in both hemispheres, which is a brain region vulnerable in old age (Vaidya et al., 2007; Insel et al., 2012). Moreover, anatomical variations in ACC morphology have been implicated in successful aging (Gefen et al., 2015). Interestingly, increased connectivity with the ACC node was largely associated with enhanced functional connectivity between the ACC and DS in aged rats. ACC-DS functional connectivity increased as a function of cognitive training in the aged rats, while the young animals showed declining ACC-DS functional connectivity following the baseline scan. In fact, declining ACC-DS connectivity was associated with improved behavioral performance, and the ability of young rats to suppress a response bias and correctly perform the WM/BAT. The ACC directly projects to the DS (Gabbott et al., 2005; Fillinger et al., 2018), and both structures are also indirectly connected through the central medial nucleus of the thalamus (Vertes et al., 2012). Given the prominent role of cholinergic and monoaminergic inputs in both regions, these data suggest that there is a dissociation between the impact of ACC activity on target neurons in the DS with age that could arise from alterations in cholinergic (Nieves-Martinez et al., 2012) and dopaminergic (Stark and Pakkenberg, 2004; Darbin, 2012) neuromodulation of the frontostriatal network. In young animals, ACC activation may serve to suppress response-based behavioral strategies leading to reduced resting state functional connectivity. In aged rats, when ACC-DS connectivity increases with training, there is no suppression of response-based strategies and behavioral performance on the WM/BAT does not improve. This idea is consistent with multiple lines of evidence. First, it is widely reported that aged animals with hippocampal-dependent spatial memory impairments tend to over utilize response-based strategies (Barnes et al., 1980; Tomas Pereira et al., 2015), that are supported by the DS (Packard and McGaugh, 1992; Graybiel, 1998; Pych et al., 2005). Second, successful performance on the WM/BAT requires animals to flexibly update their behavior based on their position in the maze. Set-shifting is compromised in aged rats (Barense et al., 2002; Beas et al., 2013; Beas et al., 2016), and this deficit has been linked to age-associated neurobiological alterations in the ACC and DS (Nicolle and Baxter, 2003; Nieves-Martinez et al., 2012). Finally, it is striking that the aged rats with less flexible and more response-driven behavior had high cellular *Arc* activity levels in the DS during performance of the WM/BAT, but not during the control spatial alternation task. It is known that neurons in DS that express *Arc* are GABAergic principal cells that are also positive for CaMKII (Vazdarjanova et al., 2006). A previous study showed that during spatial exploration, which does not evoke response-driven repetitive behavior, induces ~5% of DS neurons to express *Arc* in young rats (Vazdarjanova et al., 2006). The current study adds to our understanding of *Arc* in DS by showing that when animals employ response-driven behaviors, as the aged rats did during WM/BAT performance, there is an increased engagement of DS neurons. Taken together these multi-scale imaging data therefore suggest that engagement of the ACC during cognitive training in aged rats drives the DS to be overactive. This aberrant activity, in turn contributes to perseverative behavior and response biases that impede the ability to learn to successfully perform the WM/BAT.

An additional novel finding from the current data is the observation that rich club participation interacted with age and cognitive training. Baseline rich club participation was lower in aged compared to young rats (Figure 5a). This observation is consistent with data from human study participants that have reported less functional rich club participation in older compared to younger adults (Cao et al., 2014a). As in previous studies (Liang et al., 2018), the young rats did not show a change in rich club participation across cognitive training (Figure 5b). In contrast, the aged rats had a large increase in rich club participation between baseline and the second scan. This increase persisted in the third scan even though the aged rats displayed little to no improvements across the 27 days of cognitive testing (Figure 1c). These data are consistent with reports of network connectivity measures in humans. A previous longitudinal study showed that older adults had less network stability over time compared to young study participants (Iordan et al., 2017). It is hypothesized that rich club organization and the strength and proportion of long-distance connections plays a central role in optimizing global brain communication efficiency for normal cognition (Bullmore and Sporns, 2009, 2012; van den Heuvel et al., 2012). Presumably, at the foundation of the functional rich club are hub neurons that have long-rage projections. Interestingly, recent data have suggested that these neurons may be particularly vulnerable in advanced age with subsets of them being overrecruited during behavior in aged rats (Hernandez et al., 2018b). This over recruitment in aged animals during behavior could manifest as enhanced resting state rich club participation that ultimately reflect less adaptive networks and a reduced ability to recruit additional resources during behavior.

The notion that aged animals and older adults are less able to recruit additional resources as cognitive load increases has been formalized by the Compensation-Related Utilization of Neural Circuits hypothesis (CRUNCH). CRUNCH postulates that more neural resources are recruited by older adults during tasks with minimal cognitive load. This increased activation, could serve to compensate for a network that is compromised. As tasks become more difficult, the limits of network capacity may be reached in older adults and these compensatory mechanisms are no longer effective, leading to equivalent or less activation in older adults relative to young (Reuter-Lorenz and Cappell, 2008; Grady, 2012). The current data are consistent with CRUNCH, as the older rats showed a quick increase in rich club participation even when they were unable to correctly multi-task, suggesting that network limits in older rats are reached faster. In fact, these data along with a recent cellular imaging study showing elevated *Arc* expression in the medial prefrontal cortices of aged rats at rest (Hernandez et al., 2018b) indicate that baseline brain connectivity of older animals may be close to maximum capacity even at rest. If the capacity of a network to respond to increasing cognitive load is constrained by increased rich club participation, then aged rats may be less able to recruit additional resources while performing a difficult cognitive multi-task. Ultimately, this elevated rich club participation in old animals could contribute to the reduced dynamic range of neural activity that has been reported for older adults (Kennedy et al., 2017).

An important impact of higher rich club participation in aged rats could be to tax brain energy reserves that may already compromised in advanced age (Yoshizawa et al., 2014; Goyal et al., 2017; Hernandez et al., 2018a). Densely connected, long distance projections are metabolically costly (Bullmore and Sporns, 2012). The higher costs associated with functional connections across multiple hubs makes them particularly vulnerable to metabolic deficiencies and cellular dysfunction contributing to instability in older animals. Thus, an enticing hypothesis is that improving the metabolic capacity of older animals could restore the dynamic range and functional rich club architecture. In the future, large-scale assessment of network connectivity in conjunction with single neuron activity dynamics and metabolic function could elucidate productive therapeutic avenues for treating cognitive aging.

## Materials and Methods

### Subjects and behavioral testing

A total of 6 young (4 months old) and 6 aged (24 months old) male Fischer 344 × Brown Norway F1 (FBN) hybrid rats from the National Institute on Aging colony at Taconic Farms were used in this study. Notably, the lifespan of the FBN is greater than inbred Fisher 344 rats (Turturro et al., 1999), and many of the physical issues experienced by Fischer 344 rats are not evident in the FBN rats until they are older than 28 months (McQuail and Nicolle, 2015). Therefore, changes in performance are likely due to cognitive decline and not age-related physical impairment. Five rats in each age group were trained and scanned for the network analysis. One aged rat reached a humane endpoint prior to the *Arc* catFISH experiment and was therefore not included in this study, but this animal’s data were included in the resting state analysis. An additional young (n = 1) and aged (n = 1) rat were sacrificed directly from the home cages as a negative control to ensure that nothing unexpected occurred in the colony room on the day of the experiment to increase *Arc* expression. Expression levels in these rats was low (<5%; data not shown). Each rat was housed individually in a temperature and humidity-controlled vivarium and maintained on a reverse 12-hour light/dark cycle. All behavioral testing was performed in the dark phase.

All rats were allowed 1 week to acclimate to the housing facility prior to food restriction and initial behavioral shaping. One week after arrival, all rats were placed on restricted feeding in which 20.5 g (1.9 kcal/g) of moist chow was provided daily, and drinking water was provided *ad libitum*. Shaping began once rats reached approximately 85% of their baseline weights. Baseline weight was considered the weight at which an animal had an optimal body condition score of 3. Throughout the period of restricted feeding, rats were weighed daily, and body condition was assessed and recorded weekly to ensure a range of 2.5-3. The body condition score was assigned based on the presence of palpable fat deposits over the lumbar vertebrae and pelvic bones. Rats with a score under 2.5 were given additional food to promote weight gain. All procedures were in accordance with the National Institutes of Health Guide for the Care and Use of Laboratory Animals and approved by the Institutional Animal Care and Use Committee at the University of Florida.

Once rats reached their 85% of baseline weight, after ~1 week of restriction, they began shaping on the digital-8-shaped maze. Rats were first habituated to the testing apparatus for 10 minutes a day for 2 consecutive days, with Froot Loop pieces (Kellogg’s Company, Battle Creek, MI) scattered throughout the maze to encourage exploration. Following habituation, once rats were comfortable on the testing apparatus, they were trained to alternate between the left and right turns for 32 trials per day or 30 min. This proceeded for 1 week, then rats underwent the first baseline fMRI scan.

After the baseline scan, rats began testing on the working memory/bi-conditional association task of cognitive multi-tasking (WM/ABT; Figure 1a). During this task, rats perform continuous spatial alternations. Before returning to the center section of the maze, however, rats are ‘interrupted’ with an object discrimination problem on the choice platform in which a target object can be displaced and the animal received a Froot Loop piece. While the same object pair is presented in both the left and right choice platforms, different objects are rewarded on platform. Thus, animals must integrate information about where in the maze they are with the object information to form the correct bi-conditional association between an object and a place. On the first day of testing, objects were only partially covering the food reward for the first four trails per object (8 trials total) to encourage learning. Rats could begin with a trial turning in either the left or right direction, but on all subsequent trials, rats had to alternate turning directions. Should a rat mistakenly make a wrong turn, the trial was recorded as a working memory error and rats were not presented with the object discrimination problem. Rats were tested on this paradigm for eleven consecutive days and then given 2 days off during which the second scanning session occurred. After the scans were completed, rats testing for another 14 days and were scanned for a third and final time. After the last scan, rats were retrained on the WM/BAT with a new set of objects for the *Arc* catFISH experiment.

### Functional magnetic resonance imaging

Rats were imaged under isoflurane (1.5%) sedation (delivered in 70%N_2_/30%O_2_ at 0.1L/min). Inhalation anesthetics (e.g., isoflurane) are preferred for longitudinal fMRI experiments over intravenous injectable anesthetics due to better control over blood levels of the sedative, fast recovery and lower mortality rates in rats. Important to the present study, several studies have confirmed BOLD activation patterns at low levels of anesthesia in rats (Masamoto et al., 2007; Kannurpatti et al., 2008; Kim et al., 2010; Williams et al., 2010). Isoflurane induces dose-dependent vasodilation; thus, functional experiments must be ideally performed under doses lower than 2% (i.e., a fixed concentration between 1 and 1.5%) as was done in the current study (fixed at 1.5%). Even in human neuroimaging studies, general anesthesia (sevoflurane) did not prevent the measurement of BOLD activity and general connectivity (Riehl et al., 2018). Spontaneous breathing was monitored during MRI acquisition (SA Instruments, Stony Brook, NY). Body temperature was maintained at 37-38°C using a warm water recirculation system. A resting state fMRI dataset was collected in an 11.1 Tesla Bruker system (Magnex Scientific). The system is a Bruker AV3 HD console/Paravision 6.01 with a volume transmit (85mm inner diameter quadrature coil), and a 4-channel phase-array receive coil (Rapid Rat Phase Array). All ten rats were scanned over the span of two months in three scanning sessions: before cognitive training (first session; n_young_ = 5, n_old_ = 5), following 11 days of training (second session), and after an additional two weeks of training (third session). A 1-shot spin echo EPI sequence was acquired with acquisition parameters: TR/TE = 2000/15 ms, and 300 repetitions for a total acquisition time of 10 mins (an image was acquired every 2s), FOV = 25.6 × 25.6 mm^2^, 20 slices 1.0mm thick, and data matrix = 64 × 64. Anatomical scans for image overlay and reference-to-atlas registration were collected using a fast spin echo sequence, with the following parameters: TR/TE_eff_ = 4500/48 ms, RARE factor = 16, and number of averages = 6, FOV = 25.6 × 25.6 mm^2^, 20 slices 1.0mm thick, and data matrix = 256 × 256.

### Image processing

Brain masks were drawn manually over high-resolution anatomical scans using segmentation tools in itkSNAP (www.itksnap.org). The masks were used to crop images and remove non-brain voxels. The cropped brain images were aligned with a rat brain template using the FMRIB Software Library linear registration program *flirt* (Jenkinson et al., 2002), using previously published parameters (Colon-Perez et al., 2016). Registration matrices were saved and used to subsequently transform functional datasets into atlas space for preprocessing and analysis. Slight displacements in individual images over the series of 300 images and slice timing delays were corrected, and time series spikes were removed using Analysis of Functional NeuroImages (AFNI)(Cox, 1996). Linear and quadratic detrending, spatial blurring, and intensity normalization were also performed. Six head motion parameters and cerebroventricular and white matter signals were removed from all datasets. A voxelwise temporal band-pass filter (between 0.01 Hz and 0.1 Hz) was applied to remove brain signals that contain cardiac and respiratory frequencies.

Time series fMRI signals were extracted from each region of interest (ROI) based on the atlas-guided seed location (150 total areas, equally divided in left and right representations of each region). Signals were averaged from voxels in each ROI (Colon-Perez et al., 2016). Voxel-wise cross-correlations were carried out to create correlation coefficient (Pearson r) maps. The first 9 images in each functional time series were not used in the cross-correlation step. Pearson r maps were subjected to a voxelwise z-transformation. The two correlation maps were averaged per subject to generate a single correlation map subsequently used for statistical mapping. AFNI’s *3dClustSim* program was used to determine the adequate cluster size for a given uncorrected *p*-value. The resultant voxel cluster size at p ≤ 0.05 was used to control the level of false positive rates in the final composite statistical maps.

### Network analysis

Brain connectivity networks were analyzed using the Brain Connectivity Toolbox for Matlab (Rubinov and Sporns, 2010) and as previously reported (Colon-Perez et al., 2018; Orsini et al., 2018). Symmetrical connectivity graphs with a total 11,175 matrix entries were first organized in Matlab [graph size = n(n-1)/2, where n is the number of nodes represented in the graph or 150 ROI]. The z score values of the graphs were thresholded for each subject to create matrices with equal densities (e.g., z values in the top 15% of all possible correlation coefficients). Matrix z values were normalized by the highest z-score, such that all matrices had edge weight values ranging from 0 to 1. *Node strength* (sum of edge weights), *clustering coefficient* (the degree to which nodes cluster together in groups), *average shortest path length* (the potential for communication between pairs of structures), and *small-worldness* (the degree to which rat functional brain networks under study deviate from randomly connected networks) were calculated for these weighted graphs (Newman, 2003; Boccaletti et al., 2006; Saramaki et al., 2007). Brain networks were visualized using BrainNet (Xia et al., 2013). The 3D networks were generated with undirected edges weights *E*_undir_ ≥ 0.3. In these brain networks (or rat brain connectomes), the node size and color is scaled by the node strength and edges are scaled by z-scores.

### Tissue collection and Arc catFISH

After behavioral testing, rats were sacrificed, and tissue was collected to evaluate immediate-early gene activity during the WM/BAT and alternation tasks. Rats were placed into a bell jar containing isoflurane-saturated cotton (Abbott Laboratories, Chicago, IL, USA), separated from the animal by a wire mesh shield. Animals lost righting reflex within 30 seconds of being placed within the jar and immediately euthanized by rapid decapitation. Tissue was extracted and flash frozen in 2-methyl butane (Acros Organics, NJ, USA) chilled in a bath of dry ice with 100% ethanol (~-70°C). One additional rat in each age group were sacrificed directly from the home cage as a negative control during the experiment to ensure that disruptions within the colony room do not lead to robust nonexperimental behaviorally induced *Arc* expression. Tissue was stored at −80°C until cryosectioning and processing for fluorescence in situ hybridization.

Tissue was sliced at 20-μm thickness on a cryostat (Microm HM550) and thaw-mounted on Superfrost Plus slides (Fisher Scientific). Fluorescence in situ hybridization (FISH) for the immediate-early gene *Arc* was performed as previously described (Guzowski et al., 1999). Briefly, a commercial transcription kit and RNA labeling mix (Ambion REF #: 11277073910, Lot #: 10030660; Austin, TX) were used to generate a digoxigenin-labeled riboprobe using a plasmid template containing a 3.0 kb Arc cDNA (Steward et al., 1998). Tissue was incubated with the probe overnight, and *Arc*-positive cells were detected with antiedigoxigenin-HRP conjugate (Roche Applied Science Ref #: 11207733910, Lot #: 10520200; Penzberg, Germany). Cy3 (Cy3 Direct FISH; PerkinElmer Life Sciences, Waltham, MA) was used to visualize labeled cells, and nuclei were counterstained with DAPI (Thermo Scientific).

Z-stack images were collected by fluorescence microscopy (Keyence; Osaka, Osaka Prefecture, Japan) in increments of 1 μm. Four images (2 from superficial layers and 2 from deep layers; Figure 6a) were taken from the ACC of both the left and right hemispheres from 3 different tissue sections for a total of 24 images for each rat. Six images (3 from medial and 3 from lateral) were taken from both hemispheres of the DS for a total of 36 images per rat. The percentage and subcellular location of Arc-positive cells was determined by experimenters blind to age and order of behavioral tasks using ImageJ software with a custom written plugin for identifying and classifying cells. Nuclei that were not cutoff by the edges of the tissue and only those cells that were visible within the median 20% of the optical planes were included for counting. All nuclei were identified with the *Arc* channel off, as to not bias the counter. When the total number of cells in the z-stack were identified, the *Arc* channel was turned on to classify cells as positive for nuclear *Arc*, cytoplasmic *Arc*, both nuclear and cytoplasmic *Arc*, or negative for *Arc*. A cell was counted as *Arc* nuclear positive if 1 or 2 fluorescently labeled foci could be detected above threshold anywhere within the nucleus on at least 4 consecutive planes. A cell was counted as *Arc* cytoplasmic positive if fluorescent labeling could be detected above background surrounding at least 1/3 of the nucleus on 2 adjacent planes. Cells meeting both of these criteria were counted as *Arc* nuclear and cytoplasmic positive.

Neural activation during the WM/BAT and spatial alternation tasks was examined using the percentage of cells positive for cytoplasmic and/or nuclear Arc expression. A mean percentage of cells was calculated for each rat for each brain region and condition, so that all statistics were based on the number of animals for sample size, rather than images or cells. This avoids the caveat of inflating statistical power and having different dependent variables correlate with each other, which can be the case in nested experimental designs (Aarts et al., 2014). Critically, the order of behavior was counterbalanced across rats, with equal numbers in both age groups, such that cytoplasmic staining corresponded to OPPA task behavior and nuclear staining corresponded with alternation behavior for half of the rats and vice versa for the others. Thus, all plots showing mean percentage of *Arc*-positive cells are in reference to the task and not the epoch. Notably, as in previous studies with similar behaviors (Hernandez et al., 2018b), task order did not have a significant impact on neuron activity levels.

Potential effects of age and brain region on the percentage of cells expressing *Arc* during the different behaviors (WM/BAT versus spatial alternation) were examined with factorial ANOVAs. All analyses were performed using Statistical Package for the Social Sciences (SPSS), v.25, software. Statistical significance was considered at p values less than 0.05.

## Acknowledgments

This work was supported by National Institutes of Health National Institute on Aging grants R01 AG049722 and P50 AG047266, and the McKnight Brain Research Foundation. The authors acknowledge the support from the Advanced Magnetic Resonance Imaging and Spectroscopy (AMRIS) facility (National Science Foundation Cooperative Agreement No. DMR-1157490 and the State of Florida).

